# LXR-inducible host E3 ligase IDOL targets a human cytomegalovirus reactivation determinant

**DOI:** 10.1101/2022.11.15.516687

**Authors:** Luwanika Mlera, Donna Collins-McMillen, Sebastian Zeltzer, Jason C. Buehler, Melissa Moy, Kristen Zarrella, Katie Caviness, Louis Cicchini, David J. Tafoya, Felicia Goodrum

**Author notes:** Imanis Life Sciences, Rochester, MN 55901, USA. National Biodefense Analysis and Countermeasures Center, Fort Detrick, MD 21702, USA. PPD, Inc., 929 N Front St. Wilmington, NC 28401, USA.

## Abstract

Liver X receptor (LXR) signaling broadly restricts virus replication; however, the mechanisms of restriction are poorly defined. Here, we demonstrate that the LXR-inducible cellular E3 ligase IDOL (inducible degrader of low-density lipoprotein receptor, LDLR) targets the human cytomegalovirus (HMCV) UL136p33 protein for turnover. *UL136* encodes multiple proteins that differentially impact latency and reactivation. UL136p33 is a determinant of reactivation. UL136p33 is targeted for rapid turnover by the proteasome and its stabilization by mutation of lysine residues to arginine results in a failure to quiet replication for latency. We show that IDOL targets UL136p33 for turnover, but not the stabilized variant. IDOL is highly expressed in undifferentiated hematopoietic cells where HCMV establishes latency, but is sharply downregulated upon differentiation, a stimulus for reactivation. We hypothesize that IDOL maintains low levels of UL136p33 for the establishment of latency. Consistent with this, knockdown of IDOL impacts viral gene expression in WT HCMV infection, but not in infection where UL136p33 has been stabilized. Further, induction of LXR signaling restricts WT HCMV reactivation from latency, but does not affect replication of a recombinant virus expressing a stabilized variant of UL136p33. This work establishes the UL136p33-IDOL interaction as a key regulator of the bistable switch between latency and reactivation. It further suggests a model whereby a key viral determinant of HCMV reactivation is regulated by a host E3 ligase and acts as a sensor at the tipping point between the decision to maintain the latent state or exit latency for reactivation.

**Importance:** Herpesviruses establish life-long latent infections, which pose an important risk for disease particularly in the immunocompromised. Our work is focused on the beta-herpesvirus, human cytomegalovirus (HCMV) that latently infects the majority of the population worldwide. Defining the mechanisms by which HCMV establishes latency or reactivates from latency is important to controlling viral disease. Here, we demonstrate that the cellular inducible degrader of low-density lipoprotein receptor, IDOL, targets a HCMV determinant of reactivation for degradation. The instability of this determinant is important for the establishment of latency. This work defines a pivotal virus-host interaction that allows HCMV to sense changes in host biology to navigate decisions to establish latency or replicate.

## Introduction

Viral latency, the ability of a virus to persist indefinitely in the infected host, is a hallmark of herpesviruses. Latency is defined as a non-replicative state punctuated by sporadic reactivation events where the virus replicates productively for transmission. Human cytomegalovirus (HCMV) is a beta-herpesvirus that establishes latent infection in hematopoietic progenitor cells (HPCs) and cells of the myeloid lineage (1). Reactivation of HCMV from latency in the immunocompromised can result in life-threatening disease and is a particularly important risk factor in the context of stem cell and solid organ transplantation. The molecular mechanisms by which HCMV senses changes in the host to make decisions to maintain latency or reactivate from latency is important in developing novel strategies to control viral reactivation and subsequent CMV disease.

Latency is a complex multifactorial phenomenon dependent on viral determinants, host cell intrinsic, innate, and adaptive responses, as well as host cell signaling and chromatin remodeling (2). Current paradigms hold that HCMV reactivation is tightly connected to hematopoietic cell differentiation (3-7) or inflammatory signaling (3, 8-10). However, much remains to be understood about the precise host contributions and key viral factors that navigate the entry into or exit from latency. We and others have defined roles of *UL133-138* gene locus encoded within the UL*b*’ region of the HCMV genome for latency and reactivation. *UL138* is required for latency and the UL138 protein functions by regulating cell surface receptor trafficking and signaling (11-14), innate immune responses (15, 16) and also contributes to repression of viral gene expression (17). *UL135*, by contrast, is required for reactivation from latency, functioning in part by opposing the function of *UL138* (18-20). *UL136* is expressed as 5 protein isoforms that have differential effects on the establishment of latency or reactivation. The largest, membrane-associated isoforms, UL136p33 (UL136p33) and p26, are required for reactivation, whereas the small, soluble isoforms are required for latency in both *in vitro* and *in vivo* experimental models (21, 22). Importantly, UL136 proteins accumulate with delayed kinetics relative to UL135 and UL138 and maximal accumulation of *UL136* gene products depend on the onset of viral DNA synthesis and/or entry into late phase. Given its kinetics and dependence on viral DNA synthesis for maximal expression, we postulate that UL136 may negotiate checkpoints that function to maintain latency or commitment to reactivation from latency. Identifying host pathways regulating these viral determinants is key to defining the mechanistic basis by which HCMV senses and responds to changes in host biology to maintain latency or reactivate.

The cellular environment the virus encounters upon infection undoubtedly plays important roles in the establishment of latency with a host of restriction factors prepared to silence the viral genome, as well as a transcriptional milieu that impacts viral gene expression. As an example, we have shown that the cellular transcription factor early growth response factor-1 (EGR-1) drives the expression of UL138 for the establishment of latency (20, 23). As EGR-1 is required to maintain stemness in the bone marrow niche and is highly expressed in undifferentiated hematopoietic cells (24), cells are primed to drive the expression of UL138 for latency upon infection. However, HCMV also regulates EGR-1 expression through the differential trafficking and turnover of the host receptor tyrosine kinase, epidermal growth factor receptor (EGFR) upstream of EGR-1 (11, 19, 20) and through direct targeting by microRNAs that promote reactivation (23). Thus, HCMV has evolved a multi-pronged strategy to manipulate and utilized host pathways to regulate its gene expression, hardwiring infection into host signaling pathways. With regards to reactivation, activation of the major immediate early (MIE) transcriptional unit, encoding the major transactivators for viral replication, is highly dependent on host transcription factors in myeloid cells (25). Host transcription factors CREB, FOXO3a and AP-1 are downstream of major signaling pathways, including Src, PI3K/Akt, and MEK/ERK, and are important for re-expression of MIE genes (26-28). Further, UL7 encodes a ligand that binds the cellular Fms-like tyrosine kinase 3 receptor (Flt-3R) that when secreted upon reactivation activates PI3K/AKT and MAPK signaling to prevent cell death and stimulate differentiation for reactivation (29, 30). Through its co-evolution with the human host, the biology of HCMV infection has become tightly intertwined with the biology its host, which reflects the complex mechanisms the virus has evolved to “sense” and “respond” to changes in the cellular environment to influence “decisions’’ to maintain latency or reactivate from latency.

Here we demonstrate that the largest isoform of UL136, UL136p33, is targeted for rapid turnover relative to other UL136 protein isoforms and that its instability (t_1/2_∼1 h) is important to the establishment of latency. We identified the LXR-inducible degrader of low-density lipoprotein receptor (LDLR) IDOL (also known as myosin regulatory light chain interacting protein or MYLIP), as the host E3 ligase that targets UL136p33 for turnover. IDOL expression is induced by oxysterol activation of the liver X receptor (LXR) (31) and is regulated through hematopoietic differentiation (32, 33). In addition to LDLR, IDOL also targets VLDLR and ApoER-2 and, therefore, is an important homeostatic regulator of intracellular cholesterol (31, 34-36). The targeting of UL136p33 by IDOL suggests that the virus has evolved to be responsive to changes in cellular cholesterol in its commitment to reactivation. A model emerges whereby high levels of IDOL maintains low levels of UL136p33 for the establishment of latency, and the downregulation of IDOL by differentiation results in the accumulation of UL136p33 to drive reactivation. Indeed, induction of IDOL results in increased turnover of UL136p33 and prevents reactivation of HCMV from latency, whereas depletion of IDOL increases viral gene expression in hematopoietic cells. Our findings define UL133p33 is a key viral determinant of HCMV reactivation that acts as a sensor at the tipping point between the decision to maintain the latent state or exit latency for reactivation.

## Results

### UL136p33 is unstable and is targeted for proteasomal degradation

*UL136* is expressed as at least 5 co-terminal protein isoforms (Fig. 1A). These isoforms are encoded by transcripts with unique 5’ ends and have distinct roles in viral latency and reactivation (21, 22). The largest of these isoforms is a 33-kDa protein (UL136p33) that is required for replication in endothelial cells as well as reactivation from latency in CD34+ HPCs and in humanized mice (21). Conversely, the two small isoforms p23 and p19 suppress replication in endothelial cells and promote latency in CD34+ HPCs and humanized mice (22). The diametric nature of the UL136 system presupposes additional levels of control to coordinate and commit to one infection program over another, thereby allowing for either optimal replication/reactivation or robust latency. The UL136 isoforms, and particularly the full-length isoform UL136p33, are uniquely labile and sensitive to mechanical lysis while being processed (22). Based on this, we hypothesized that the concentration of some UL136 isoforms is regulated by increased proteolysis, and that the instability of UL136p33 may be important to establishing or maintaining latency.

**Figure 1.**
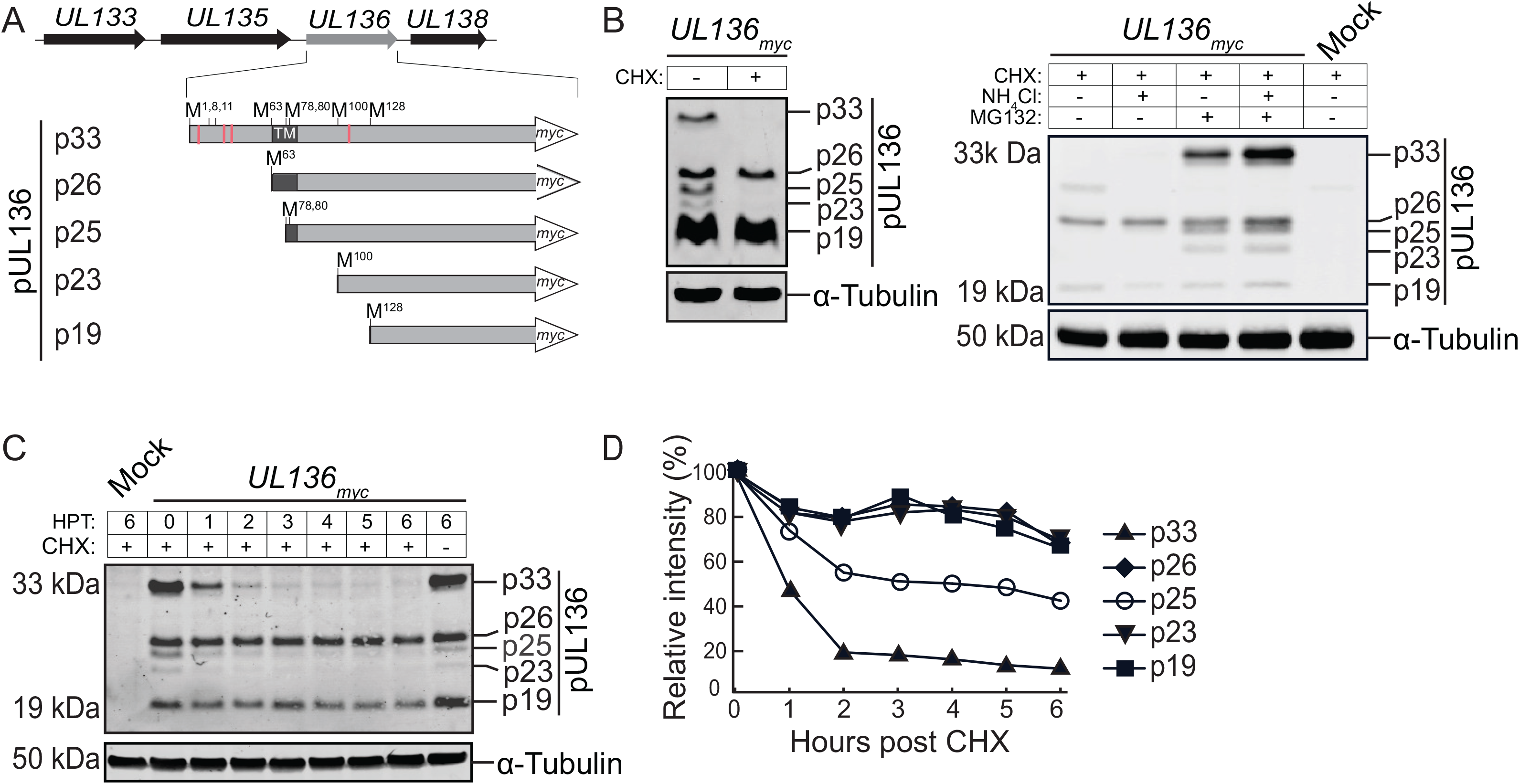
UL136p33 isoform is rapidly turned over by the proteasome. (A) Schematic depiction of the *UL133-UL138* polycistronic gene locus and UL136 protein isoforms. A C-terminal myc tag is used to detect UL136 isoforms. Red lines represent lysine residues, and the black box shows the putative transmembrane domain. (B-D) Fibroblasts were infected with TB40/E-*UL136_myc_* virus at a MOI of 1. (B) At 48 hpi, cells were treated with 50 µg/ml cycloheximide (CHX) for 6 h. Alternatively, mock- or TB40/E-infected cells were treated with CHX in the context of proteasomal (MG132) or lysosomal (NH_4_Cl) inhibition (right panel). UL136_myc_ protein isoforms or tubulin as a loading control were detected by immunoblotting using by a monoclonal α-myc antibody or tubulin, respectively (left panel). (C) Cells were untreated or treated with CHX at 48 hpi and UL136 decay was analyzed over a 6-hour time course by immunoblotting. (D) Relative quantification of UL136 isoforms normalized to tubulin. HPT, hours post treatment

To explore the possibility that UL136 isoforms have different stabilities, we analyzed the levels of UL136 isoforms in infected fibroblasts treated with cycloheximide (CHX) to halt protein synthesis. Cycloheximide treatment resulted in a striking loss of UL136p33, p25, and p23, but not p26 or p19 (Fig. 1B, left panel). These results indicated differential stability whereby UL136p33, p25, and p23 may be subject to more rapid turnover, while other UL136 isoforms are relatively stable. Proteasome inhibition robustly rescued UL136p33, p25, and p23 isoforms (Fig. 1B, right panel). Inhibition of the lysosome with NH_4_Cl had no effect on rescue of any of the UL136 isoforms; however, inhibition of both the lysosome and proteasome showed a somewhat additive effect. Together, these results indicate that the rapid turnover of UL136p33, p25, and p23 is regulated by the proteasome.

To establish the kinetics of isoform turnover, we analyzed the decay of UL136 isoforms in fibroblasts infected with TB40/E-*UL136*_myc_ and treated with CHX at peak UL136 isoform expression (Fig 1C). UL136p33 decayed with a half-life of 1 h (t_1/2_=1 h) and p25 exhibited a t_1/2_=3 h, while all other isoforms were stable over the 6 h time course (Fig 1D). The p23 isoform accumulates to very low levels making the decay of this protein is difficult to determine. The rapid decay of UL136p33 relative to the other isoforms, as well as its significance in HCMV reactivation prompted us to focus on determining the factor(s) contributing to its instability.

### Lysine to arginine mutations stabilize UL136p33

UL136 has four lysine (K) residues (denoted by red lines in Fig. 1A), three of which lie upstream of the transmembrane domain (TMD) and are unique to the p33 isoform. To determine if the turnover of UL136p33 was regulated by ubiquitin-mediated proteolysis, we substituted all lysine residues for arginine (R) through site-directed mutagenesis in an expression vector encoding UL136p33 to generate *UL136p33_myc_ΔK→R*. The p33_ΔK*→*R_ mutant had a t_1/2_=5 h, compared to ∼1 h for UL136p33 following CHX treatment (Fig. 2A). This observation, together with the stabilization of UL136p33 by proteasomal inhibition (Fig. 1B), suggests that UL136p33 turnover is regulated by ubiquitin-mediated proteasomal decay. Further, the rapid turnover of exogenously expressed UL136p33 (Fig. 2A, left panel) indicates that UL136p33’s degradation is not dependent on virus or virus-induced factors.

**Figure 2.**
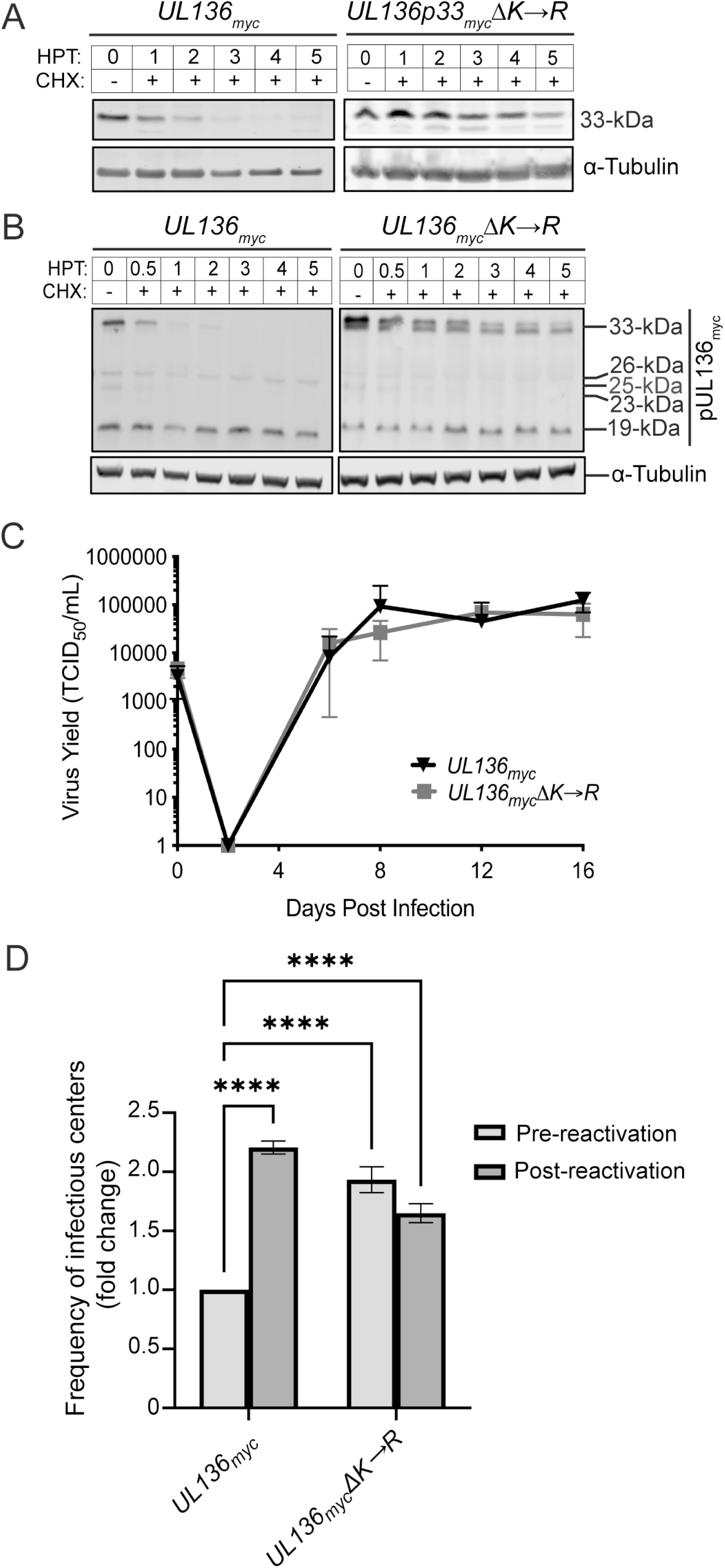
Stabilizing UL136p33 results in virus that fails to establish latency. (A) Fibroblasts were transduced with lentiviruses engineered to express p33_myc_ or p33_K→R_ and protein decay was analyzed by immunoblotting following CHX treatment. (B) Fibroblasts were infected with TB40/E-*UL136_myc_* or -*UL136_myc_ΔK→R* viruses (MOI =1) and treated with CHX at 48 hpi or left untreated. Protein decay was analyzed as described in Figure 1C. (C) Fibroblasts were infected at a MOI of 0.02 and samples collected for TCID_50_ at indicated times. All data points are an average of 3 biological replicates and error bars represent standard deviation. (D) CD34+ HPCs were infected with *UL136_myc_* or *UL136_myc_ΔK→R* viruses (MOI=2). Infected cells were purified by FACS at 1 dpi seeded into long term culture. At 10 days, half of the culture was reactivated by seeding intact cells by limiting dilution into co-culture with fibroblasts in a cytokine-rich media to promote myeloid differentiation (reactivation). The other half of the culture was mechanically lysed and seeded onto fibroblasts as the pre-reactivation control. The frequency of infectious centers was determined for each sample by extreme limiting dilution. (C-D) The data represents replicates with cells from 3 independent donors. The means were compared with two-way analysis of variance (ANOVA) with a Tukey post hoc test and the were no significant differences. **, *p*<0.01; ***, *p*<0.001.

To determine the importance of UL136p33 instability in infection, we introduced the lysine to arginine mutations into the *UL136_myc_* gene within the bacterial artificial chromosome clone of TB40/E (*UL136_myc_ΔK→R*). We measured the relative rates of decay in fibroblasts infected with either the parental *UL136*_myc_ or *UL136_myc_ΔK→R* virus and treated with CHX and observed that p33_ΔK→R_ was stabilized during infection relative to wild type UL136p33 (Fig. 2B). Interestingly, we stabilization of UL136p33 diminished levels of UL136p26 and p25 isoforms by mechanisms that have yet to be determined.

We then performed multistep growth curves on fibroblasts infected with *UL136_myc_* or *UL136_myc_ΔK→R* to analyze the impact of stabilizing UL136p33 on virus replication. No change in the kinetics or yield of viral growth was observed during the productive infection in fibroblasts when UL136p33 is stabilized relative to the WT infection (Fig. 2C). This was not surprising since the UL136p33 isoform, while important for replication in endothelial cells and CD34+ HPCs, is dispensable for replication in fibroblasts (21, 22). In summary, UL136p33 instability is the result of lysine-dependent, proteasomal degradation and mutating lysine residues to arginine in *UL136* stabilizes the protein with no effect on virus replication in fibroblasts.

Since UL136p33 is required for reactivation from latency in CD34+ HPCs (21), we hypothesized that the instability of UL136p33 allows for a tunable concentration gradient where low levels are important for the establishment of latency and accumulation past a threshold is important for reactivation. To test this hypothesis, we performed a latency assay in primary human CD34+ HPCs infected with the parental *UL136*_myc_ or *UL136_myc_ΔK*→*R* viruses. *UL136_myc_*-infected CD34+ HPCs generated >2-fold frequencies of infectious centers following reactivation compared to the pre-reactivation control (Fig. 2D). However, the pre- and post-reactivation frequencies of infectious centers were not different in the *UL136_myc_ΔK→R* infection and were almost two-fold above *UL136*_myc_ pre-reactivation. These results indicate that HCMV replication in CD34+ HPCs is not quieted when UL136p33 is stabilized, and that this virus does not establish a latent infection. In a separate study, we have shown that the stabilization of UL136p33 results in increased virus replication (a failure to establish latency) in a humanized mouse model of latency (37). As is the case with many viral proteins, UL136 isoforms, even with a myc epitope tag, are not expressed to detectable levels in hematopoietic cells, and we have been unsuccessful in generating antibodies to UL136. Nevertheless, given the requirement for UL136p33 for reactivation and replication in CD34+ HPCs, the *UL136_myc_ΔK→R* infection phenotype is consistent with accumulation of UL136p33 beyond a threshold critical for commitment to replication.

### The host E3 ligase IDOL targets UL136p33 for proteasomal degradation

Ubiquitination of cargo by E3 ligases is an important step in targeting a protein for proteasomal degradation (38, 39). Because UL136p33 is rapidly turned over by the proteasome even in the absence of infection and lysine to arginine substitutions stabilized UL136p33, we hypothesized that a host E3 ligase targets UL136p33 for turnover. We identified UL136 interacting proteins by a yeast two-hybrid screen (Y2H) using the C-terminal region of UL136 (amino acids 141-240) as bait. Intriguingly, we identified the host E3 ligase IDOL (**i**nducible **d**egrader **o**f **l**ow-density lipoprotein receptor), also known as MYLIP, as a candidate host interactor of UL136 (Supporting information, SI Table 1). IDOL induces the degradation of the low-density lipoprotein receptor family members LDLR, VLDLR, and ApoER-2 (35, 36). The degradation of these receptors is a physiological response to increased cholesterol, which is modified to oxysterols that activate the LXR transcription factor to induce *IDOL* expression (36, 40, 41).

**Table 1.** Antibodies used in this study.

| Antibodies |  |  |
| --- | --- | --- |
| Rabbit monoclonal anti-ApoER-2 | Abcam | Cat#ab108208 |
| Rabbit polyclonal anti-FLAG tag | Sigma-Aldrich | Cat#F7425 |
| Rabbit monoclonal anti-GAPDH (clone D16H11) | Cell Signaling Technology | Cat#5174 |
| Rabbit polyclonal anti-HA tag | Sigma-Aldrich | Cat#H6908 |
| Mouse HDAC1 (clone 10E2) | Santa Cruz Biotech | Cat#sc-81598 |
| IE1/2 (3H4) | Princeton University/Dr. Tom Shenk | N/A |
| Rabbit monoclonal anti-Myc tag (clone 71D10) | Cell Signaling Technology | Cat#2278 |
| Tubulin (clone DM1A) |  | Cat#T6199 |
| UL135 | Open Biosystems | Custom antibody |
| Goat anti-mouse secondary antibody (IR Dye 680) | LI-COR Biosciences | Cat#926-68070 |
| Goat anti-rabbit secondary antibody (IR Dye 800) | LI-COR Biosciences | Cat#926-32211 |

To determine if IDOL targets UL136p33, we analyzed the ability of IDOL to interact with and impact UL136p33 levels. We validated the interaction between IDOL and UL136p33 by immunoprecipitating (IP) UL136p33-IDOL complexes from cells transiently overexpressing myc-tagged UL136p33 or HA-tagged IDOL. Transient overexpression is necessary due to the lack of commercially available antibodies specific and sensitive enough to detect or immunoprecipitate endogenous IDOL. Pulldown of IDOL_HA_ or p33_myc_ coprecipitated the alternant partner (Fig. 3A), confirming the interaction between UL136p33 and IDOL. Further analysis indicated that co-expression of IDOL with UL136p33 resulted in a ∼60% reduction in UL136p33 levels compared to when UL136p33 was co-expressed with an empty vector (Fig. 3B top and quantified in Fig. 3C), consistent with IDOL driving UL136p33 turnover. By contrast, p33_ΔK→R_ levels were not affected by IDOL co-expression (Fig. 3B bottom and quantified in Fig. 3C), in agreement with the hypothesis that ubiquitination on lysine residues is required for UL136p33 turnover via the proteasome. Taken together, IDOL functions as a host E3 ligase that targets UL136p33 for proteasomal degradation.

**Figure 3.**
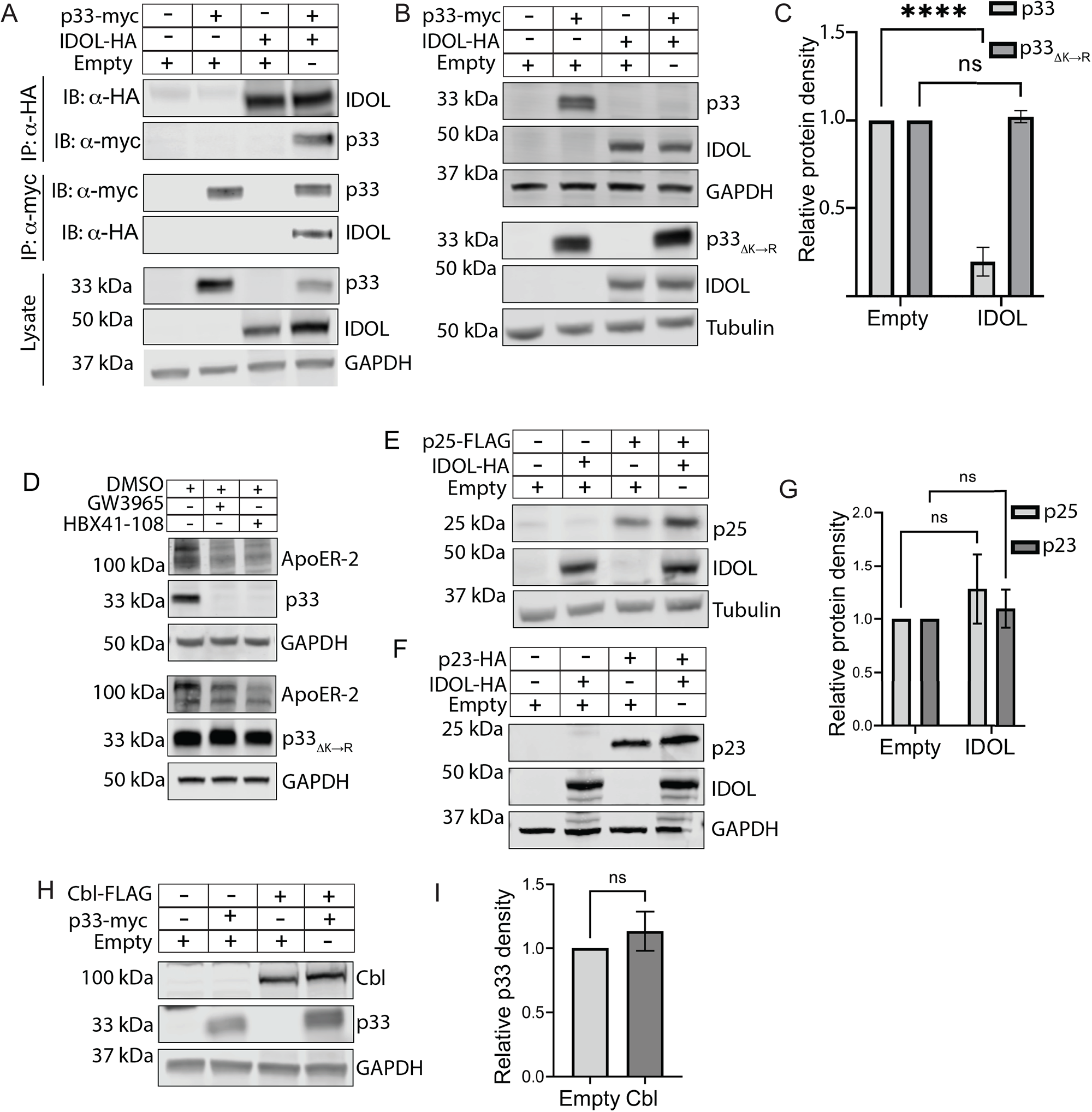
The host E3 ligase IDOL specifically targets UL136p33 for turnover. (A) HEK cells were transiently transfected with plasmids expressing UL136p33, IDOL, empty vector, or the indicated combinations. UL136p33 or IDOL was immunoprecipitated from protein lysates using α-myc or α-HA magnetic beads. Protein was detected by immunoblotting using the antibodies indicated. Lysates were blotted as a control for expression and GAPDH used as a loading control. (B) HEK cells were transiently transfected with plasmids expressing p33_ΔK→R_, IDOL, or empty vector. Proteins were detected by immunoblotting using α-myc or α-HA monoclonal antibodies. (C) Protein densities of UL136p33 (from panel A) or p33_ΔK→R_ (from panel B) relative to their loading control. (D) HEK cells were transiently transfected to express UL136p33 or p33_ΔK→R._ IDOL was induced 48 hours later with 2 µM of LXR-agonists, GW3965 or HBX41-108, or vehicle (DMSO) for 6 h. Detection of ApoER-2 was used as a proxy for validating the induction of IDOL. p33_myc_, p33_K→R_, and tubulin as a loading control were detected by immunoblotting using α-myc or α-tubulin. (E-G) HEK 293T cells were transfected with plasmids expressing (E) IDOL, p25 or empty vector or (F) IDOL, p23 or empty vector. Proteins detected by immunoblotting using antibodies to FLAG or HA to detect the indicated proteins. Tubulin or GAPDH were used as loading controls for normalization. (G) Relative p25 or p23 densities. (H) HEK 293T cells were co-transfected with plasmids expressing UL136p33, the E3 ligase Cbl or empty vector. Proteins were detected by immunoblotting using antibodies to the myc epitope tag or cbl. (I) Relative UL136p33 densities were measured and quantified. All data points represent the mean of at least 3 independent replicates and the means were compared by two-way ANOVA with a Tukey post hoc test. Error bars represent standard deviation. ns, non-significant *p*>0.05; ****, p<0.0001.

We next induced endogenous IDOL expression with an LXR agonist GW3965 or LXR-independent agonist HBX41-108 in cells transiently expressing UL136p33 or p33_ΔK→R_. Because endogenous IDOL cannot be detected by western blot using available antibodies (34, 42, 43), the induction of IDOL was verified by the loss of its target ApoER-2 (Fig. 3D). Induction of IDOL, by both GW3965 and HBX41-108 resulted in a reduction of UL136p33 (Fig. 3D, top), but not p33_ΔK→R_ (Fig. 3D, bottom). These results further confirm that endogenous IDOL targets UL136p33 for ubiquitin-dependent degradation.

The Y2H bait was composed of the C-terminal region of UL136 spanning amino acids 141-240 downstream of the predicted transmembrane domain. These residues are shared by all the UL136 isoforms (Fig. 1A). Further, our data indicated that p25 and p23 proteins are also labile following CHX treatment and these smaller isoforms could be rescued, like UL136p33, by inhibiting the proteasome (Fig. 1B, right panel). The p25 and p23 isoforms have a single lysine which is located downstream of the TMD in the full length UL136 isoform (Fig. 1A). Transient co-expression of IDOL with either p25 or p23 did not lead to any notable turnover of these proteins (Fig. 3E-G). These data collectively led us to conclude that IDOL specifically targets UL136p33 for turnover, and it does not mediate the degradation of p25 and p23. The mechanism by which p25 and p23 are turned over remains to be elucidated and could reflect differential trafficking and localization of the proteins (22).

To further define the specificity of IDOL-mediated turnover of UL136p33, we co-expressed UL136p33 with another E3 ligase, Cbl. Cbl targets tyrosine kinase receptors, such as EGFR, for ubiquitination and subsequent degradation by the lysosome (44). Levels of UL136p33 were unaffected by co-expression with Cbl (Fig. 3H-I), indicating some specificity of IDOL for proteasomal turnover of UL136p33.

### IDOL impacts UL136p33 protein levels in HCMV-infected primary fibroblasts

In the context of a productive infection in fibroblasts, we found that infection with either the parental *UL136_myc_* or *UL136_myc_ΔK→R* induced *IDOL* transcripts by at least 5-fold compared to mock-infection by reverse transcriptase, quantitative PCR (RT-qPCR), (Fig. 4A). *IDOL* transcript upregulation was sustained over time during infection and peaked at close to 10-fold by 3 dpi. No changes in *IDOL* transcript levels were detected in mock controls over the same time-course. Because *IDOL* induction is a consequence of increased cellular cholesterol, we measured cholesterol content in infected cells to correlate it to the upregulation of *IDOL* transcripts. Using this approach, we noted that HCMV increased cholesterol concentration by at least 29% from 1 dpi onward (Fig. 4B), consistent with previous findings on HCMV-modulation of lipid biosynthesis (45-50). Taken together, infection in fibroblasts upregulates expression levels of IDOL transcripts, which is correlated with increased intracellular cholesterol levels.

**Figure 4.**
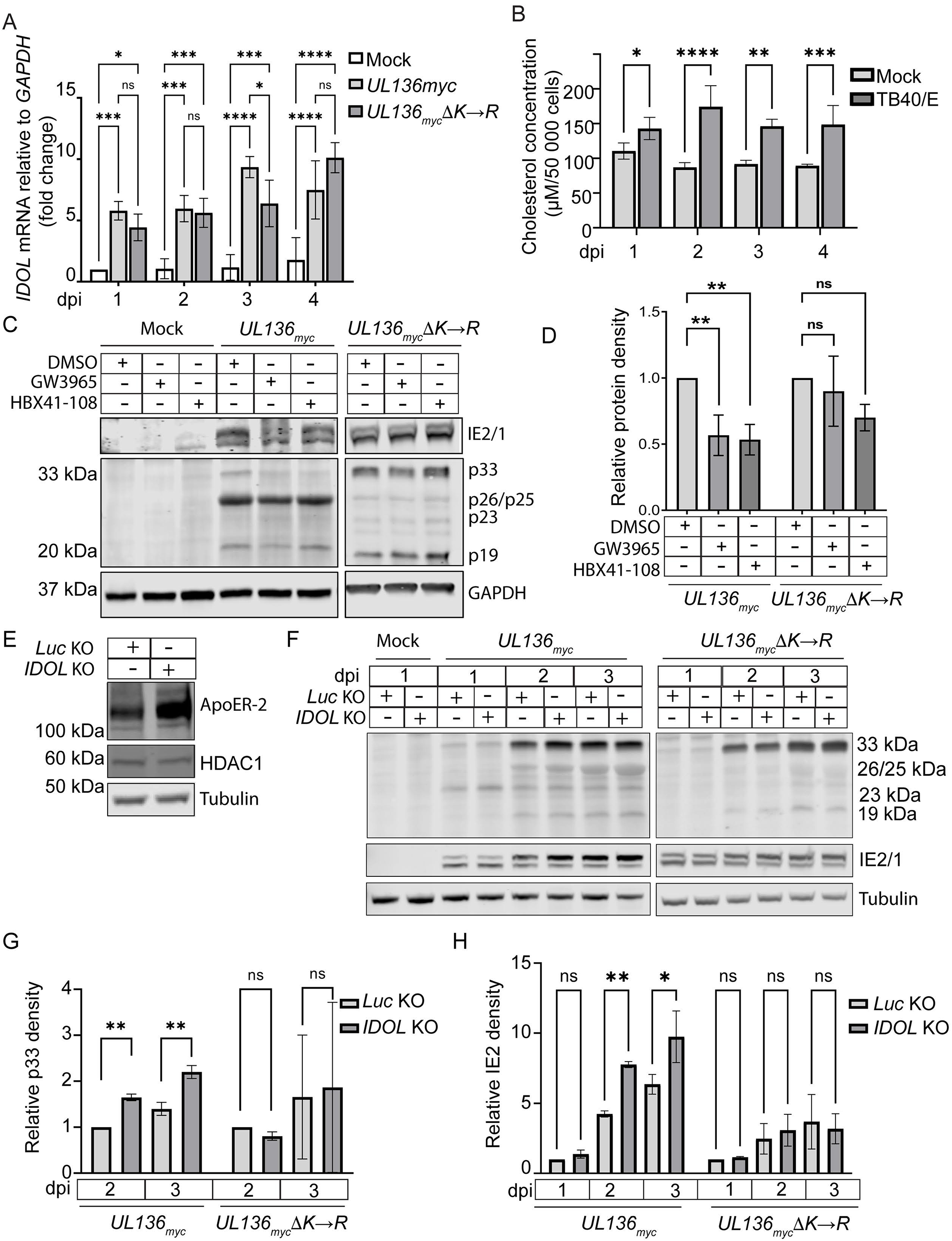
IDOL expression in HCMV-infected fibroblasts impacts UL136p33 stability. (A) Fibroblasts were mock-infected or infected with *UL136_myc_* or *UL136_myc_ΔK→R* viruses at a MOI of 1 and total RNA extracted at indicated time points. RT-qPCR was used to quantify relative *IDOL* transcripts using the ΔΔCT method. *IDOL* transcripts were normalized to GAPDH and compared to expression levels in mock samples at 1 dpi. (B) Fibroblasts were infected with TB40/E at a MOI of 1 followed by incubation in serum-free media. Cholesterol concentrations were measured and compared to mock-infected cells at 1 dpi. (C) Fibroblasts were mock-infected or infected with *UL136_myc_* or *UL136_myc_ΔK→R* viruses at a MOI of 1. At 2 dpi, IDOL expression was induced by treatment with 2 µM GW3965 or HBX41-108 for 6 h followed by western blot evaluation of UL136p33 levels (left). Relative UL136p33 levels are quantified in (D). (E) *IDOL* KO cell lysates were analyzed by western blot to evaluate ApoER-2 expression levels compared to a *Luc* KO control. To evaluate CRISPR/CAS9 off-target effects we measured HDAC1 in *Luc* or *IDOL* KO cell lysates. (F) *UL136_myc_* or *UL136_myc_ΔK→R* viruses were used to infect either *Luc* or *IDOL* KO cells at a MOI of 1. Cell lysates were collected at indicated times and evaluated for UL136p33. IE1 and IE2 protein levels that are quantified in panels G and H. (G) Quantification p33 protein densities relative to the Luc KO at 2 dpi in each infection condition, *UL136_myc_* and *UL136_myc_ΔK→R*. (H) Quantification of IE2 protein densities relative to *Luc* KO at 2 dpi in each infection condition, *UL136_myc_* and *UL136_myc_ΔK→R*. Quantitative data points represent the mean of 3 independent replicates and error bars represent standard deviation. Differences in the means were compared using two-way ANOVA with a Tukey post hoc test. ns, p>0.05; *, p<0.05; **, p<0.01; ***, p<0.001; ****, p<0.0001.

To address the effect of HCMV-mediated induction of IDOL on UL136p33 levels in a productive infection, we again induced IDOL expression with the LXR agonist GW3965 or HBX41-108 in TB40/E-*UL136*_myc_-or -*UL136*_myc_ΔK→R-infected fibroblasts. IDOL induction resulted in a 50% reduction in the levels of UL136p33 but did not significantly affect p33_ΔK→R_ levels (Fig. 4C and D). Diminished levels of smaller UL136 isoforms are again apparent in the *UL136*_myc_ΔK→R infection. We next disrupted *IDOL* expression in fibroblasts to answer the question whether IDOL depletion would rescue UL136p33 turnover. CRISPR-mediated knockout (KO) of *IDOL* using guide RNAs to disrupt all the isoforms of IDOL was validated by Sanger sequencing and increased ApoER-2 as a proxy for *IDOL* KO (Fig. 4E). Further, CRISPR KO of *IDOL* had no effect on HDAC1, a potential off-target hit when knocking out *IDOL*. In fibroblasts depleted of IDOL, UL136p33 levels increased at 2 and 3 dpi compared to luciferase KO controls, but did not affect levels of p33_ΔK→R_ (Fig. 4F and quantified in 4G). Further, IE2 levels were also significantly increased in IDOL KO cells infected with *UL136_myc_*, but not changed by IDOL KO in the *UL136_myc_ΔK→R* infection (Fig. 4H). Together, these results indicate that IDOL regulates UL136p33 concentration, which indirectly impacts IE expression in the context of infection.

### LXR induction and IDOL restricts HCMV gene expression and reactivation in models of latency

HCMV latency is studied in a variety of models. Due to limitations inherent to each model, there is no perfect model and, therefore, we explore latency questions using multiple complementary models. To explore the role of IDOL in regulating UL136p33 levels in latency and reactivation, we have used two models of HCMV: (1) THP-1 cell line model and (2) CD34+ primary HPC model. In the THP-1 monocytic cell line model, cells are infected in an undifferentiated state for the establishment of latency. Viral gene expression is detected early (1 dpi) following infection but is silenced as the virus establishes a latent-like state (51). Re-expression of viral genes for reactivation is stimulated by differentiating the cells into macrophages with a phorbol ester, tetradecanoylphorbol acetate (TPA) (52-57). This is a strong model for looking at transcriptional changes associated with latency and reactivation; however, this model is limited in that THP-1 cells do not robustly or consistently support viral genome synthesis or production of viral progeny following TPA induction, a requirement for bona fide reactivation. The gold-standard for HCMV latency is the ability of the virus to establish and reactivate from latency in primary human CD34+ HPCs (58). In contrast to THP-1 cells, the primary CD34+ HPC model supports viral genome synthesis and the formation of infectious progeny in response to reactivation. However, the model is limited by substantial donor-variability associated with primary human cells, heterogeneity of the population, the availability of cells in numbers to provide sufficient starting material for many assays, and the ability to manipulate these cells (e.g. gene knockdowns or overexpression) without affecting their differentiation or quantities sufficient for HCMV infection assays. Further, because reactivation of HCMV from CD34+ HPCs occurs at low frequency (1 in ∼9,000 cells), analysis of molecular events associated with reactivation is challenging. The use of both models yields complementary insights into the patterns of gene expression upon infection or reactivation (THP-1) and virus production (CD34+ HPCs) to understand latency and reactivation.

Dynamic gene expression changes are associated with monocyte-to-macrophage differentiation (32, 59, 60). Because IDOL is highly expressed in bone marrow (61), we hypothesized that *IDOL* expression may be dictated by the differentiation state of the cell and would regulate UL136 levels in a differentiation-dependent manner. Indeed, we found that *IDOL* was highly expressed in undifferentiated THP-1 monocytes prior to infection but was sharply downregulated upon THP-1 cell differentiation (Fig. 5A). Further, while no significant differences in *IDOL* expression were observed between uninfected cells and *UL136_my_*_c_ infection, *UL136_myc_ΔK→R* infection significantly reduced *IDOL* expression in the undifferentiated, latent phase (Fig. 5A). This result suggests that UL136p33 levels may feedback to impact IDOL expression in a cell type-dependent manner.

**Figure 5.**
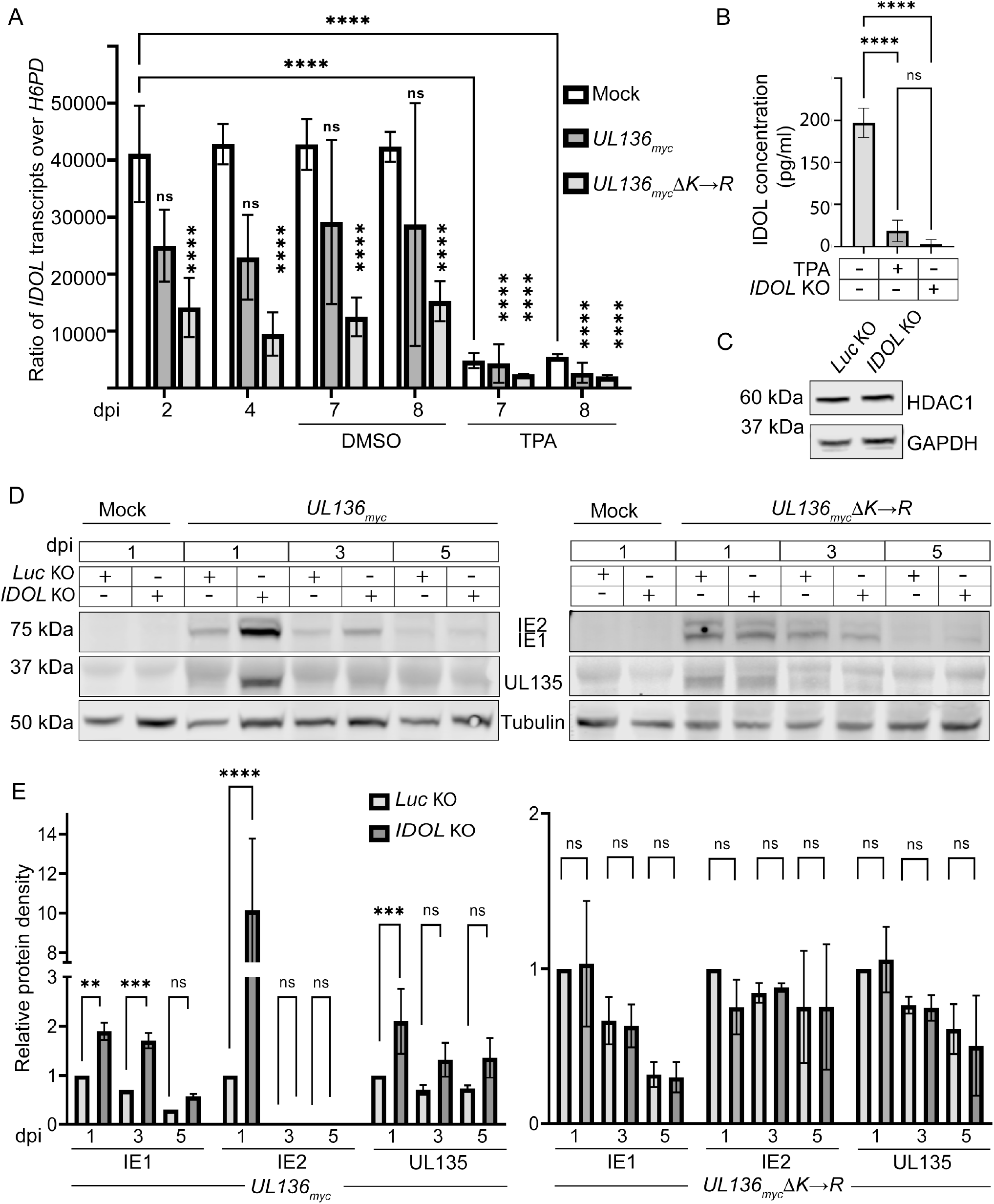
IDOL impacts viral protein expression in a THP-1 cell latency model. (A) Total RNA was extracted from mock-, *UL136_myc_*- or *UL136_myc_ΔK→R-*infected THP-1 cells at indicated time points. At 5 dpi, cells were treated with TPA to induce differentiation or the vehicle control (DMSO). cDNA was synthesized from the RNA and qPCR was used to determine the relative expression levels of *IDOL* transcripts compared to the low copy cellular housekeeping gene *H6PD* using the ΔΔCT method. Validation of *IDOL* KO in undifferentiated THP-1 cells was confirmed by ELISA. Where indicated, cells were treated with TPA as a positive control for the loss of IDOL. Immunoblot of HDAC1 in *Luc* control or *IDOL* KO THP-1 cells to rule out off-target CRISPR/CAS9 effects. (D) *IDOL* or *Luc* KO THP-1 cells were infected with TB40/E-*UL136*_myc_ (top panel) or *-UL136_myc_ΔK→R* (bottom panel) at a MOI of 2 and cell lysates were collected at indicated time points. IE1/2 and UL135 accumulation was evaluated by immunoblotting protein lysates over a time course using a monoclonal α-IE1/2 and polyclonal α-UL135. (E) IE1, IE2 and UL135 were quantified, and levels normalized to Tubulin and are shown relative to levels present at 1 dpi in the *Luc* KO control for *UL136_myc_* and *UL136_myc_ΔK→R* infections. Note that comparisons are made between *Luc* KO and *IDOL* KO, not between *UL136_myc_* and *UL136_myc_ΔK→R* infections. All data points are representative as the mean 3 independent replicates and error bars represent standard deviation. The means were compared by two-way ANOVA with a Tukey post hoc test. ns, p>0.05; *, p<0.05; **, p<0.01; ****, p<0.0001.

To determine how IDOL impacts HCMV infection in hematopoietic cells, we knocked out *IDOL* in THP-1 cells using CRISPR. The knockdown was validated using an ELISA assay for the detection of IDOL protein (Fig. 5B); IDOL protein was reduced in undifferentiated cells to levels induced by differentiation. Further, *IDOL* KO had no off-target effects on HDAC1 (Fig. 5C). Given the challenges of detecting UL136 isoforms, particularly in hematopoietic cells, we examined IE1, IE2 and UL135 gene expression as proxies for determining the effect of *IDOL* KO on infection in THP-1 cells. UL135 and IE1/2 proteins are detected early during infection in THP-1 cells but diminish as the virus establishes latency between 3 and 5 dpi (Fig. 5D-E) (52). *IDOL* KO resulted in an increased initial burst in the expression of IE1, IE2, and UL135 at 1 dpi relative to control *Luc* KO cells in *UL136*_myc_ infection. Notably, the major HCMV transactivator, IE2, was detected in *UL136_myc_* infection in the IDOL KO condition, but not in the *Luc* KO control. The increased viral gene expression in the IDOL KO is consistent with increased viral activity. However, gene expression was still relatively quieted by 5 dpi, suggesting that induction of IDOL does not fully prevent the silencing of viral gene expression in this system and that other viral or cellular factors are required. As we have never identified a condition or mutant virus that is not ultimately silenced in THP-1 cells, this is not unexpected and it may reflect the robust program for restriction and silencing in THP-1, indicated by the inability of these cells to support viral DNA synthesis and progeny formation even with TPA treatment. UL135 and IE1, as well as IE2, were detected in *UL136_myc_ΔK→R* infection of THP-1 cells. Importantly, *IDOL* KO had no effect on IE1, IE2, or UL135 expression in *UL136_myc_ΔK→R* infection (Fig. 5D-E), consistent with a role for IDOL in regulating the establishment of latency by targeting UL136p33 for turnover. In contrast to *UL136*_myc_ infection, IE2 was detected in all conditions of the *UL136_myc_ΔK→R* infection and is not increased further by IDOL KO. While a potent transactivator, IE2 feeds back to negatively regulate the major immediate early promoter and its own expression (62-65). The differences in IE2 expression in IDOL KO in *UL136*_myc_ and *UL136_myc_ΔK→R* infection may reflect complexities in the regulation of IE gene expression or the possible dysregulation and loss of middle UL136 isoforms in *UL136_myc_ΔK→R* infection (Fig. 2). Further work is required to understand this relationship. These results indicate that loss of IDOL increases viral gene expression in *UL136_myc_* infection at least at early times following infection of THP-1 cells, suggesting a role for IDOL in regulating HCMV infection in undifferentiated THP-1 cells. This finding reflects IDOL-induced changes in UL136p33 since the stabilized variant does not differentially express genes in an IDOL-dependent manner. This finding is consistent with a role for IDOL in driving instability of UL136p33 and the role of UL136p33 in driving HCMV replication in hematopoietic cells (66). Further, these data indicate that UL136p33 may negatively regulate IDOL expression in hematopoietic cells, as the stabilized form of UL136p33 results in greater repression of IDOL transcripts by HCMV infection.

To further explore the importance of IDOL in latency, we turned to the primary CD34+ HPC experimental model (58). IDOL is expressed in undifferentiated CD34+ HPCs and, similar to our findings in the THP-1 cell model, its expression is downregulated by differentiation (Fig. 6A). Further, the LXR-agonist GW3965 induces IDOL expression in either infected or uninfected CD34+ HPCs (Fig. 6B). To measure the impact of LXR induction on reactivation in primary CD34+ HPCs, we isolated CD34+ HPCs infected with TB40/E-*UL136*_myc_ or -*UL136*_myc_*Δ*K→R (GFP+) by fluorescent activated cell sorting and seeded the cells into long-term bone marrow cultures. To analyze infectious virus production, infected CD34+ HPC cultures were split at 10 dpi. Half the cells were seeded by limiting dilution into co-culture with fibroblasts and cytokine-rich media to stimulate reactivation from latency. The other half was mechanically lysed and lysates were seeded by limiting dilution onto fibroblast monolayers to determine the virus present prior to reactivation (pre-reactivation). Relative to the DMSO control, GW3965 treatment restricted *UL136*_myc_ reactivation (Fig. 6C), consistent with a requirement for accumulation of UL136p33 for reactivation. By contrast, *UL136_myc_ΔK→R* did not establish latency and LXR induction, and presumably increased levels of IDOL, did not restrict replication of *UL136_myc_ΔK→R.* Given the limited availability of human CD34+ HPCs, data from two independent donors are shown for figure 6C. These results are consistent with the model that IDOL restricts virus reactivation in a manner dependent on its ability to turnover UL136p33 since LXR induction has not effect of the infection the *UL136_myc_ΔK→R*. Taken together with the THP-1 experiments, these results suggest that IDOL-driven UL136p33 instability is required for latency and that induction of IDOL suppresses reactivation, presumably through suppressing the accumulation of UL136p33 (Fig. 7).

**Figure 6.**
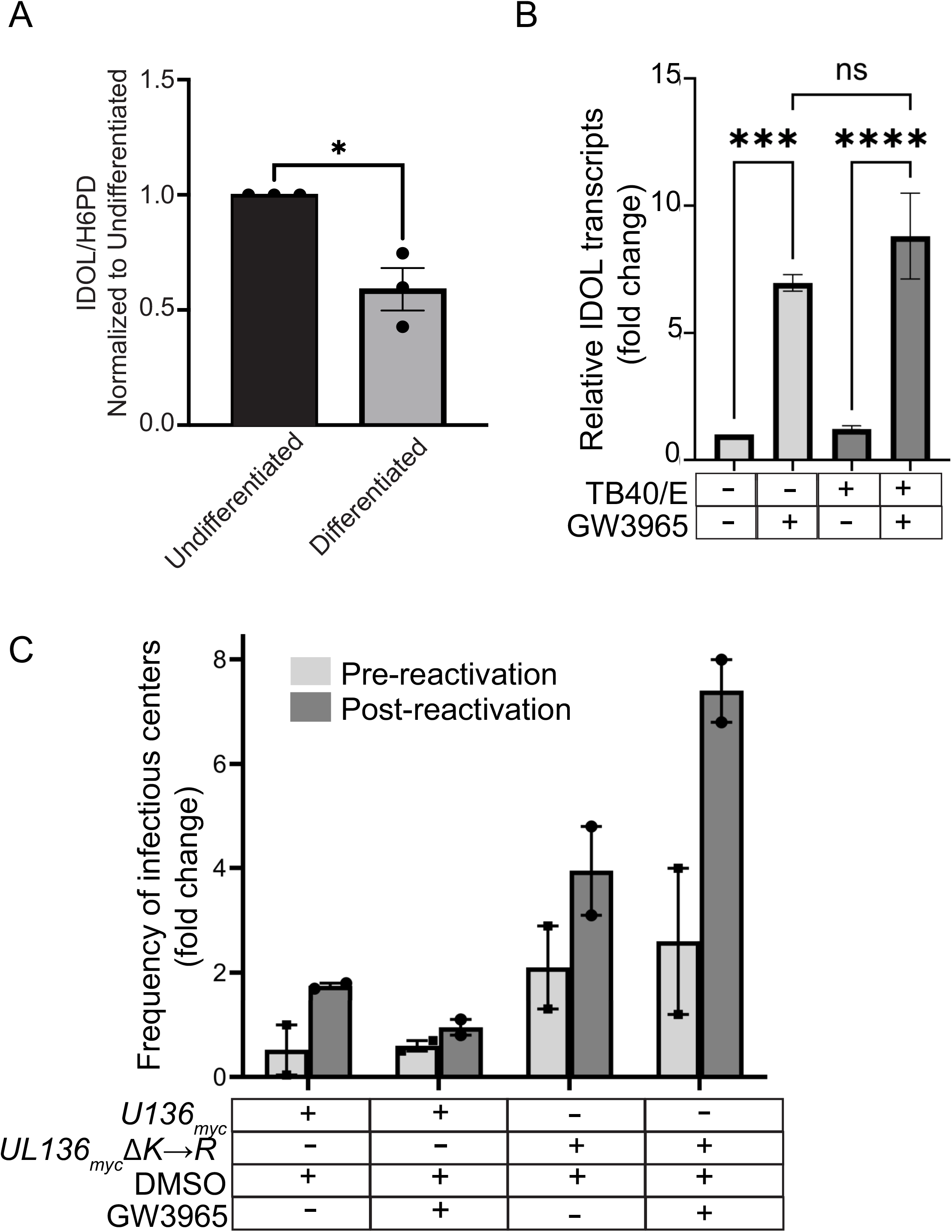
Induction of IDOL in CD34+ HPCs inhibits reactivation. (A) CD34+ HPCs were cultured for 5 days in media with cytokines to stimulate differentiation or left undifferentiated. RNA was isolated and transcripts encoding IDOL were quantified by RT-qPCR relative to the H6PD housekeeping gene using the ΔΔCT method. Values were normalized to the undifferentiated population and graphed for statistical analysis. Statistical significance was calculated using unpaired student t test and represented by asterisks. Graphs represent the mean of three replicates and error bars represent SEM. (B) CD34+ HPCs were mock- or TB40/E-infected at a MOI of 2. At 1 dpi, cells were treated with 1 µM GW3965 for 6 h and total RNA isolated for qPCR to measure *IDOL* transcript levels relative to *H6PD*. The data points represent the mean from 3 independent experiments. The means were analyzed by two-way ANOVA with a Tukey post-hoc test. ***p<0.001; ****, p<0.0001. (C) CD34+ HPCs were infected at a MOI of 2. At 24 hpi, infected CD34+ cells were isolated by FACS and seeded into long term culture. GW3965 (1 μM) was added to induce IDOL at the time of reactivation. At 10 days, half of the culture was reactivated by seeding intact cells by limiting dilution into co-culture with fibroblasts in a cytokine-rich media to promote myeloid differentiation (reactivation). The other half of the culture was mechanically lysed and seeded onto fibroblasts as the pre-reactivation control. The frequency of infectious centers was determined for each sample by ELDA. The data points represent replicates from 2 independent donors and the range is indicated by squares (pre-reactivation) and dots (post-reactivation).

**Figure 7.**
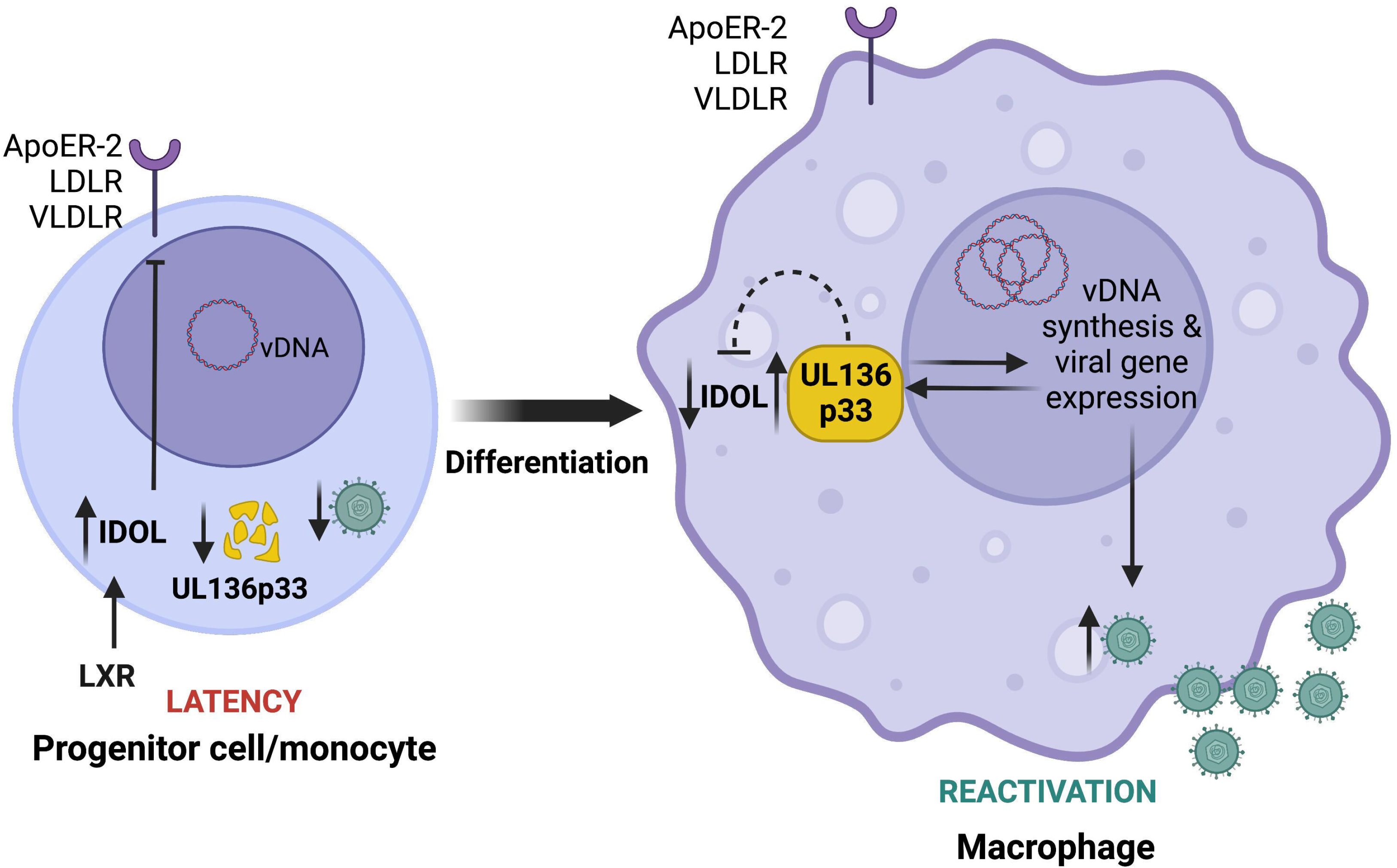
Model for differentiation-dependent IDOL regulation to control UL136p33 levels. Progenitor cells are relatively small and intolerant of high cholesterol, which they control, in part, through the expression of IDOL to reduce cholesterol/lipid uptake. We propose that high levels of IDOL maintain low levels of UL136p33 to restrict HCMV replication for the establishment of latency following infection. Upon cell differentiation, the downregulation of IDOL is observed and is expected to correspond to increased accumulation of UL136p33 to drive reactivation and replication, consistent with our finding that depletion of IDOL increases HCMV gene expression and that an LXR agnonist restricts reactivation. We have previously shown that UL136p33 levels further accumulate in late phase upon the commitment to viral DNA synthesis. Therefore, following reactivation, we postulate that UL136p33 levels increase due to a combination of decreased turnover and increased synthesis. Further, because stabilization of UL136p33 results in corresponding downregulation of IDOL expression, UL136p33 may feedback to suppress *IDOL* gene expression. This work implicates IDOL as a host factor that fine-tunes levels of UL136p33 to control decisions to enter and exit latency. UL136p33 acts as a sensor at the tipping point between the decision to maintain the latent state or exit latency for reactivation where its instability is critical for the establishment or latency and its accumulation critical for reactivation. This study suggests that LXR signaling and cholesterol homeostasis may impact HCMV latency and reactivation and will be explored in future studies. Figure created with Biorender.com.

## Discussion

Latent viruses require mechanisms to sense the cellular environment, filter noise and respond to host cues to maintain a quiescent latent state or to reactivate replication (1). While the decision to enter or exit latency is often depicted as a simple binary on/off switch, the reality is more likely that a number of successive checkpoints must be crossed, each requiring thresholds in viral or cellular activities to be reached in making the commitment to productive reactivation. Developing strategies to control entry into or exit from HCMV latency and ultimately control CMV disease rests upon identifying the virus-host interactions that govern the decisions around latency and reactivation. Here we demonstrate the existence of an unstable viral determinant of reactivation, UL136p33 (Fig. 1) and that its instability is important for the establishment of latency in CD34+ HPCs (Fig. 2). We demonstrate that its instability is dependent upon a host E3 ligase, IDOL, which is regulated by hematopoietic differentiation (Fig. 3). Knockdown of IDOL levels increases viral gene expression (Fig. 4) and LXR induction restricts replication in hematopoietic models of latency (Fig. 5-6). Our results support a model, rich in hypotheses for further investigation, whereby UL136p33 is maintained at low levels for the establishment of latency because IDOL levels are high (Fig. 7). Upon differentiation, IDOL levels fall (Fig. 5A and 6A), allowing UL136p33 levels to rise, which facilitates HCMV crossing at least one threshold towards reactivation. UL136p33 accumulation is amplified and re-enforced by the onset of viral DNA synthesis and entry into late phase (22) and high levels of UL136p33 further suppresses IDOL expression (Fig. 5A). These findings define novel virus-host interactions that govern entry into and exit from latency. Moreover, these findings suggest that by utilizing IDOL to regulate levels of a critical viral determinant for reactivation, the virus evolved to hardwire this decision into LXR signaling, which ultimately controls IDOL levels. Future work will define the mechanisms by which UL136 regulates HCMV infection and commitment to replication in hematopoietic cells. Understanding the relationship between UL136 isoforms, to other *UL133-UL138* locus-encoded proteins and to host interactors will be critical to mechanistic insights.

*IDOL* is transcriptionally upregulated by excess cholesterol derivatives (oxysterols) that are sensed by LXR and in a cell type-dependent manner (36, 67-70). RNA sequencing to classify the tissue-specific expression of genes across a representative set of all major human organs and tissues revealed that IDOL is highly expressed in the bone marrow, where CD34+ HPCs and monocytes are generated (71). The level of IDOL expression in the BM is only second to the placenta. High expression of IDOL in undifferentiated progenitor cells (Fig. 6A) or in monocytes (Fig. 5A) suggests they are intolerant of high cholesterol levels. Consistent with this, undifferentiated THP-1 monocytes express significantly higher levels of cholesterol efflux genes, such as CES1, and express significantly lower levels of influx genes (72). By contrast, THP-1 monocyte-derived macrophages upregulate cholesterol influx genes, such as scavenger receptor A1 (SR-A1) and CD36 to become lipid-loaded macrophages or foam cells. Further, statin treatment to lower cholesterol reduces CD34+ HPC populations in the circulation and high cholesterol promotes CD34+ HPC proliferation, myelopoiesis as well as monocyte and granulocytic differentiation (73-75). While carefully designed studies are required to understand how statins specifically affect HCMV latency and/or reactivation, these findings suggest the possibility that sterol homeostasis impacts HCMV latency and reactivation. Untreated hypercholesterolemia might lead to increased hematopoietic differentiation and a higher frequency of HCMV reactivation events, whereas statin treatments may restrict HCMV reactivation and replication. Given the role demonstrated here for IDOL in the regulation of cholesterol and UL136p33 expression, the high levels of IDOL expression make progenitor cells an ideal environment for HCMV latency in which UL136p33 is suppressed by rapid turnover. It is intriguing to speculate a possible role of UL136p33 as a viral sensor of changes in LXR signaling compatible with reactivation.

HCMV infection remodels the cellular lipid metabolic landscape by upregulating *de novo* cholesterol biosynthesis during a productive infection (46, 49, 50). It is, therefore, expected that HCMV infection would induce IDOL expression during a replicative infection. While IDOL and cholesterol are induced by infection in fibroblasts (SI Fig. 2A-B), IDOL expression was not induced by infection in hematopoietic cells. In fact, IDOL expression was downregulated by *UL136*_myc_*ΔK→R* infection (Fig. 5A). This disparity in the regulation of IDOL likely reflects cell type-specific differences, including tolerance for cholesterol and differences in the nature of the infection in each cell. While cholesterol increases in macrophage cells, they also have higher tolerance to cholesterol than HPCs or monocytes (76-78). Accordingly, despite increased cholesterol in macrophages, IDOL levels are strikingly downregulated in THP-1 cells induced to differentiate with TPA (Fig. 5A) or CD34+ HPCs are induced to differentiate with cytokines (Fig. 6A).

IDOL recognizes a conserved FERM domain (WxxKNxxSI/MxF) within the cytoplasmic tail of its targets (ApoER-2, LDLR and VLDLR), to mediate ubiquitination for lysosomal degradation (31, 35, 36, 79). Canonical IDOL targets reside at the plasma membrane and are typically internalized via the endocytosis and trafficked to the lysosome (79). By contrast, UL136p33 is predominantly Golgi-associated (22), lacks a FERM domain, and is degraded via the proteasome (Fig. 1B). Differences in localization and trafficking pathways between natural IDOL targets and UL136p33 may differentially direct the degradation pathway. It remains to be determined how IDOL recognized and targets UL136p33, but it is possible that the two proteins interact indirectly.

Activation of LXR in human foreskin fibroblasts with GW3965 prior to HCMV infection has been shown to depress expression levels of viral proteins, such as IE2, pp28, pp65 and gB (80). GW3965 treatment is also reported to induce IFN-γ (81), a cytokine which directly inhibits IE1/2 mRNA expression (82). These findings suggest the possibility that the effects of LXR agonists on UL136p33 levels are due to a global repression of viral gene expression. However, our results argue against this scenario. First, we observed no effect on IE gene expression at 2 dpi in infected fibroblasts treated with GW3965 or HBX41-108 (Fig. 4C-D), consistent with the attenuation of IFN-γ at later times (83). Second, specific depletion of *IDOL* increases UL136p33, consistent with IDOL-specific effects on UL136p33 levels. Nevertheless, to avoid this potential pitfall, we treated cells in our experiments with these agonists following the establishment of infection and IE gene expression in our studies.

The LXR axis is important for the biology of other herpesviruses. A recent report demonstrates that the alpha herpesvirus pseudorabies virus (PRV), etiologic agent of the economically important Aujeszky’s disease, has evolved to inhibit LXR at both transcription and protein levels (84). Using an inverse LXR agonist SR9243, Wang and colleagues showed that inhibiting LXR increased PRV replication, whereas 3 different LXR agonists and the oxysterol 22R-hydroxycholesterol all inhibited PRV replication (84). However, PRV can be rescued from the suppressive effects of an LXR agonist by supplementing with cholesterol (84), consistent with a viral need for cholesterol to replicate. LXR also inhibits the murine gamma herpesvirus MHV68 as it replicates better in *LXR^-/-^* macrophages and, contrary to PRV, MHV68 infection increases LXRα (85). Importantly, LXRα restricts reactivation of MHV86 in peritoneal cells, but not in splenocytes (86). The cell type-dependent role of LXR signaling for reactivation suggests the possibility that a LXR signaling might control a MHV68 determinant of reactivation in peritoneal cells that is dispensable in splenocytes. Similarly, disruption of or stabilization of UL136p33 has no impact on replication in fibroblasts (Fig. 2C) (22), but strongly impacts replication in HPCs (Fig. 2D) and endothelial cells (21). Viral factors encoded by MHV68 and regulated by LXR signaling have yet to be determined.

In addition to herpesviruses, LXR signaling suppresses the replication of other viruses, including Hepatitis C Virus (HCV) (87, 88), human immunodeficiency virus (HIV) (89-91), and Newcastle disease virus (NDV) (92). Intriguingly, activation of LXR signaling suppresses hepatitis C virus (HCV) replication in an IDOL-dependent manner. The suppression of HCV is presumably due to the requirement of LDLR for an entry receptor since once infection is established, IDOL induction does not seem to have an effect on HCV replication (93). However, there are mixed reports on the requirement for LDLR for HCV entry or its specific function in HCV replication (94-96).

While many host pathways are implicated in the regulation of herpesvirus latency, the interplay between viral determinants and their regulation by host mechanisms to terminate activity is less well appreciated, particularly for HCMV. The expression of HSV-1 ICP0, a critical protein for reactivation and replication, is suppressed by the host neuronal microRNA-138 to facilitate latency (97, 98). Moreover, ICP0 was recently shown to be regulated by the host E3 ligase, TRIM23, which is in turn, countered by the HSV-1 late protein, γ_1_34.5 (99). While speculative, TRIM23 may moderate ICP0 protein levels to suppress reactivation, where γ_1_34.5 ensures the commitment to replication by blocking TRIM23-mediated turnover of ICP0. Further, the E3 ubiquitin ligase, MDM2, restricts levels of the Kaposi’s sarcoma-associated herpesvirus (KSHV) transactivator, RTA/ORF50, acting as pro-viral factor for latency (100). RTA/ORF50 in turn induces degradation of vFLIP through the host Itch E3 ubiquitin ligase to attenuate vFLIP-driven NFκB signaling for reactivation (101, 102). Coopting and regulating host E3 ligases to target viral proteins for latency allows infections to respond quickly to host cues (e.g., prior to changes requiring transcription of genes and translation of proteins) for reactivation. Further, it will be important to determine if UL136p33 feeds back to regulate *IDOL* expression based on the observation that stabilization of UL136p33 represses *IDOL* expression in undifferentiated THP-1 cells (Fig. 5A). It will be important to further define the mechanisms by which UL136p33 interacts with host pathways to impact viral gene expression and promote context-dependent replication and reactivation.

The evolution of viruses to depend on host factor-mediated turnover of critical viral determinants is shared across highly divergent systems. Bacteriophages establish a dormant lysogenic state to cope with their variable and unpredictable environment (103). Bacteriophage λ lysogeny depends on the competition between repressors and activators of gene expression that serve as a gatekeeper for the lytic pathways (104, 105). As a primary example, competitive binding of the lysogeny determinant CI and the immediate early gene Cro at regulatory operator elements puts CI-Cro at the heart of a bistable switch controlling lysogenic-lytic decisions. Another determinant, CII, is expressed with later kinetics and is also critical to the commitment to lysogeny (106, 107). CII promotes expression of CI while it represses expression of the anti-terminator Q, that must accumulate to a threshold to drive lytic gene expression. Critical effectors in bacteriophage lysogeny, such as CI and CII, are susceptible to destruction by host proteases that are induced in response to stresses, such as DNA damage (108, 109). Similar to the model emerging for UL136-IDOL interaction in regulating a determinant of reactivation for latency, the decision to establish lysogeny depends on the accumulation of lysogenic effectors and is responsive to host determinants regulated by environmental cues. Further, while CI is robust, the expression of CII, like UL136, is sensitive to genome copy number or other fluctuations in the system, providing additional checkpoints in regulating the decision.

With respect to the system emerging for HCMV, the *UL133-UL138* locus coordinates the expression of multiple genes impacting infection fate decisions. While UL135 and UL138 are not known to directly regulate gene expression, their opposing impact on host EGFR/PI3K signaling (11, 18, 20) has parallels to the Cro-CI bistable genetic switch. Further, UL136p33 is expressed with later kinetics than the UL135/UL138 determinants, and robust UL136p33 expression is dependent on crossing a threshold of commitment to viral DNA synthesis (22). In addition, UL136p33 is remarkably unstable compared to other *UL133-UL138* genes and other UL136 isoforms, and, as we have shown here, its instability is dependent upon a host E3 ligase regulated by differentiation-linked changes in IDOL expression. Although the UL136p33 protein is undetectable in THP-1 monocytes or hematopoietic cells, KO of *IDOL* increases expression of genes critical for replication (e.g., IEs and UL135) and stabilization of UL136p33 results in a virus that replicates in the absence of a stimulus for reactivation in vitro (Fig. 2D), but also in a humanized mouse model of latency (37). While UL135 is critical for overcoming the repressive effects of UL138 for successful reactivation, we assert that the subsequent accumulation of UL136p33 beyond a threshold is also important for commitment to the replicative state (Fig. 7). It is important to understand the relationship between UL135 and UL136p33 in the commitment to HCMV reactivation (37) since as both are required for reactivation and how latency-promoting UL136p23/19 isoforms (21) may contribute to the establishment of latency or counter the progression towards reactivation to maintain a latent infection. This is a particularly intriguing question as the stabilization of UL136p33 not only affects levels of IDOL mRNA, but also affects levels of other UL136 isoforms (Fig. 2B). Finally, these findings indicate that the LXR-IDOL axis may be a potential target for controlling HCMV reactivation clinically.

## Methods

### Yeast two-hybrid screen

A yeast screen for human cellular interacting proteins of pUL136 was performed using the Matchmaker Gold Yeast Two-Hybrid System (Clontech) and the Mate and Plate Universal Human Library (Clontech). This yeast two-hybrid universal library was constructed from human cDNA that has been normalized to remove high copy number cDNAs (overrepresented transcripts) to facilitate the discovery of low copy number novel protein-protein interactions. All protocols were carried out according to the manufacturer’s guidelines. Yeast dual transformation and yeast mating was performed according to established protocols from Clontech. Yeast prey vectors were isolated from Y187 yeast and subsequently transformed into DH10B *E. coli*. After selection for *E coli* containing yeast prey vectors, the plasmids were isolated and sequenced to identify interacting partners. Candidate interactors are listed on SI Table 1.

### Cells and viruses

Human primary embryonic fibroblasts (MRC-5; ATCC) and human kidney embryonic 293T (HEK 293T; ATCC) cells were maintained in Dulbecco’s Essential Media (DMEM). The DMEM was supplemented with 10% fetal bovine serum (FBS), 10 mM HEPES, 1 mM sodium pyruvate, 2 mM l-alanyl-glutamine, 0.1 mM nonessential amino acids, 100 U/ml penicillin, and 100 μg/ml streptomycin. CD34+ hematopoietic progenitor cells (HPCs) were isolated from de-identified medical waste following bone marrow isolations from healthy donors for clinical procedures at the Banner-University Medical Center at the University of Arizona (58). THP-1 cells were maintained in RPMI1640 (Gibco) supplemented with 10% FBS, 1 mM sodium pyruvate, 0.1 mM β-mercapto-ethanol (Sigma).

The wild type TB40E/05 strain expressing GFP as a marker for infection (110) was engineered to fuse the myc epitope tag in frame at the 3’ end of *UL136* (*UL136_myc_* virus) using BAC recombineering as described previously (22). The *UL136_myc_ΔK→R* virus was constructed following methods previously described (22). For generation of *UL136*_myc_ΔK→R, *UL136_myc_ΔK→R* pGEM-T Easy construct was created by site-directed Phusion mutagenesis according to the manufacturer’s instruction (New England Biolabs). Inverse PCR was used to mutagenize lysines 4, 20, 25, and 113 to arginine using the primers: 5’-GACGTTGGAAATAGATGGCGGCGTCGAAGGCCCTGAGTCGC-3’ (forward); CAAGTCCCACGTCATTTCTGGCATCTCCACGCCCCTGACTGACAT-3’ (reverse) for the first 3 lysine residues in the N-terminal region and 4^th^ lysine was mutagenized to arginine with the forward primer 5’-ACGAGCGGCAGAGAAGCAGACGGC-3’, and reverse primer GGTGCCGACGGCACCTCTCAGGATAATG-3’. Following verification by Sanger sequencing, a region of the *UL136K-Rmyc* pGEM-T Easy was PCR amplified from bp 854 of UL135 [UL135(bp854)] to UL138 (bp139). This PCR product was gel purified and recombined with a ΔUL136<GALK> BAC as previously described (22). BAC integrity was tested by enzyme digest fragment analysis and sequencing of the UL133/8 viral genomic region. All BAC genomes were maintained in Escherichia coli SW102, and viral stocks were propagated by transfecting 15 to 20 μg of each BAC genome, along with 2 μg of a plasmid encoding UL82 (pp71), into 5 × 106 MRC-5 fibroblasts and subsequently purified and stored as previously described (22).

### Immunoblotting

Cells were washed twice in PBS (Gibco) and lifted from cell culture vessels with 0.25X trypsin (Gibco). Cells were washed in cold PBS, pelleted at 1000 xg for 2 minutes followed by storage at -80 °C before lysis in RIPA buffer. Total protein was quantified using a BCA assay and 50 μg of each sample was prepared for loading onto 12% precast PAGE gels (GenScript). Wet transfers were performed onto 0.45 uM pore-sized PVDF membranes (Immobilon). Blocking was achieved by 5% skimmed milk in 1X TBST, or 5% BSA in 1X TBST. Primary and secondary antibodies used for western blot analyses are indicated in Table 1.

### RT-qPCR assays

Cells were washed twice in cold PBS and lysed in 400 μl of RNA/DNA lysis buffer (Zymo Research). Total cellular RNA was extracted according to the manufacturer’s protocol for Quick-DNA/RNA minipreps. cDNA was synthesized from 1 μg of DNAse-treated RNA using the Vilo cDNA synthesis kit (Invitrogen, Thermo Fisher Scientific). qPCR reactions were prepared with *IDOL* TaqMan assay (ID Hs00982312_m1; ThermoFisher Scientific), Platinum^TM^ Quantitative PCR SuperMix-UDG w/ROX (ThermoFisher Scientific) and 50 ng cDNA. qPCR reactions were run in a QuantStudio 3 real time PCR machine (Applied Biosystems). Gene expression was normalized to GAPDH (Hs01922876_u1) or H6PD (Hs00188728_m1) and the ΔΔCT method was used to calculate relative gene expression levels (111, 112). Alternatively, cDNA was quantified using LightCycler SYBR Mix kit (Roche) and corresponding primers to IDOL (BIORAD PrimePCR^TM^ Primers) and H6PD (Forward: 5’-GGACCATTACTTAGGCAAGCA-3’ Reverse: 5’-CACGGTCTCTTTCATGATGATCT-3’). Assays performed on Light Cycler 480 and corresponding software. Relative expression was determined using the ΔΔCT method normalized to H6PD.

### Co-immunoprecipitation

Cells were washed twice with PBS and trypsinized. The PBS-washed cell pellet was lysed on ice for 30 minutes in a lysis buffer made of 100 mM Tris/HCl (pH 7.4), 100 mM NaCl, 50 mM NaFl, and 0.5% NP-40. The complete protease inhibitor cocktail (Roche) was added to the lysis buffer according to manufacturer recommendations. Lysates were clarified by centrifugation at 3000 x g for 5 minutes at 4 °C. A milligram of the lysate was incubated with Pierce anti-myc or anti-HA magnetic beads for 1 h at 4 °C. The beads were washed 3 times and protein eluted according to manufacturer’s protocol.

### CRISPR/CAS9-mediated knock out of *IDOL*

Guide RNAs (gRNAs) used to knock out IDOL were designed using online gRNA design tools at atum.bio and synthego.com. Sequences of the gRNAs are shown in Table 2. The gRNAs were cloned into lentiCRISPR v2 (113). Lentiviruses to deliver the gRNAs were generated by co-transfecting the lentiCRISPR v2 constructs with pDMG.2 and psPAX2 in HEK 293T cells. MRC-5 and THP-1 cells were transduced at a MOI of 3 and 10, respectively. Transduced cells were selected with 2 μg/ml puromycin for 2 days. Validation of knock-out was done by blotting for ApoER-2, Sanger sequencing or quantifying levels of IDOL by ELISA. The Synthego gRNA design algorithm (synthego.com) indicates potential off-target genes for the gRNAs and we selected *HDAC1* to evaluate potential off-target effects by western blot analyses.

**Table 2.**
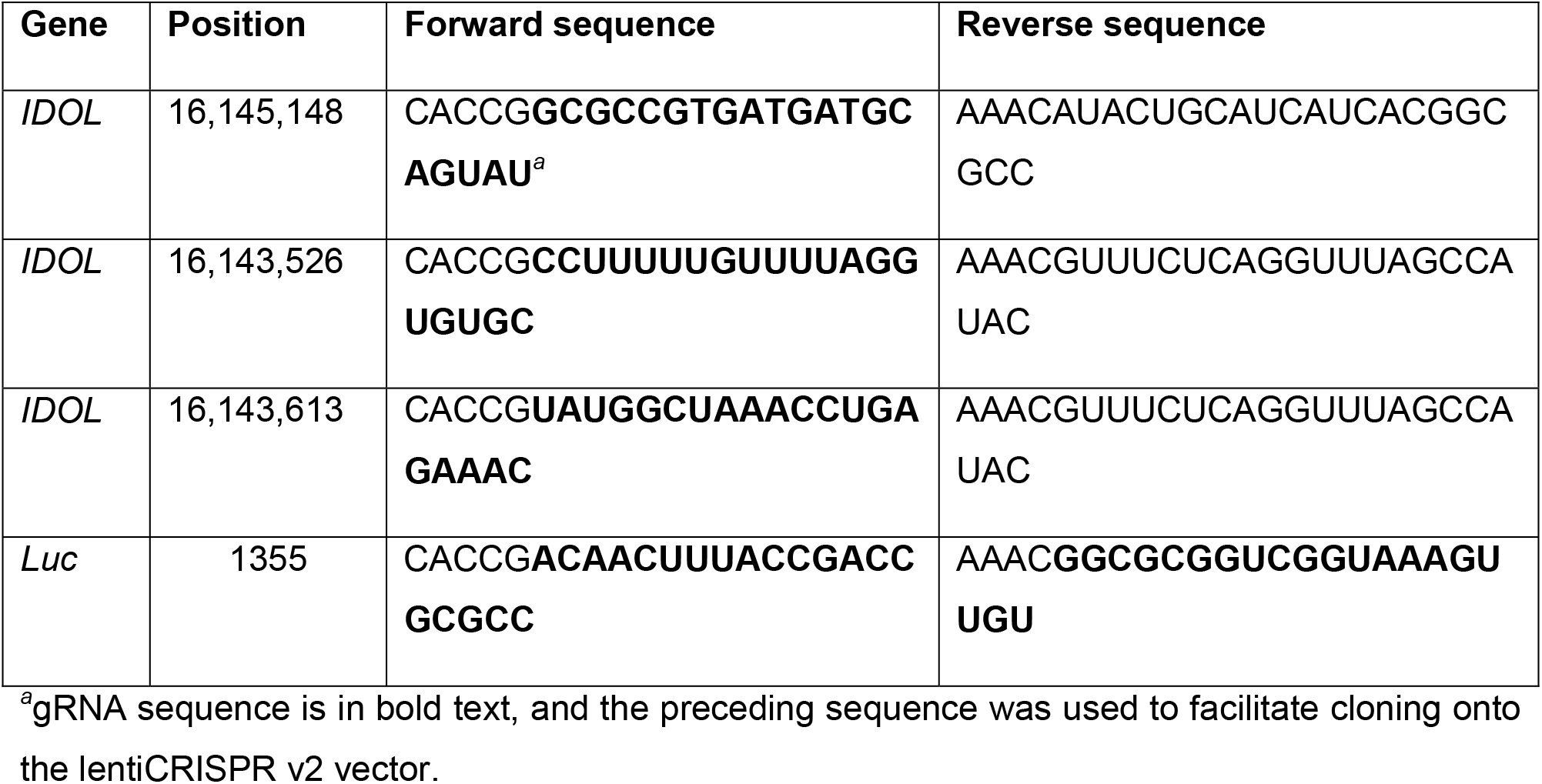
gRNAs used to KO *IDOL* and *luciferase (Luc)* gene (from *Photinus pyralis*)

### IDOL ELISA

We used a sandwich ELISA assay (Cusabio) to quantify IDOL protein levels in THP-1 cells. Cells were washed in cold PBS and resuspended at 10^8^ cells/mL followed by freezing at -20 °C. Following 3 freeze-thaw cycles, IDOL was quantified according to the manufacturer’s protocol. Optical densities were determined using a Clariostar plate reader (BMG Labtech) and sample values were interpolated from a standard curve in GraphPad Prism v9.

### Cholesterol assay

Fibroblasts were seeded at 3 x 10^5^ cells per well in 6-well dishes and incubated overnight. Infections were done at a MOI of 1 for 1 h and the inoculum was removed followed by washing cells twice with PBS. Infected cells were incubated in serum-free media and 50 000 cells were harvested at appropriate time points. Samples were prepared using a cholesterol Glo bioluminescent assay (Promega) according to the manufacturer’s instructions. Relative light units were measured using a Clariostar Plus plate reader (BMG Labtech) and sample values were extrapolated from a standard curve in GraphPad Prism v9.

### Latency assays

Latency assays in CD34+ HPCs were carried out as described previously (58). IDOL expression was induced prior to reactivation by treatment with 1 μM GW3965. Infections for latency in THP-1 cells were done as described previously (52)

## Acknowledgements

This study was supported by the National Institutes of Allergy and Infectious Diseases/ National Institutes of Health (NIAID/NIH) grants AI079059 and AI131598 to FG. KZ was supported by a training grant (T32 AG058503) from the National Institute of Aging. We are grateful for the support of the Flow Cytometry and Human Immune Monitoring Shared Resource, supported through the Cancer Center Support Grant P30 CA023074 awarded to University of Arizona. We appreciate technical assistance from Lincoln Gay. We thank Drs. James Alwine (University of Pennsylvania), Lynn Enquist (Princeton University), Jean Wilson (University of Arizona), and John Purdy (University of Arizona) for critical discussions on this work.

## Notes

### Competing Interest Statement

The authors have declared no competing interest.

### Summary of Updates

Updated in response to review; data figures added.

