## Supplementary material for "LXR-inducible host E3 ligase IDOL targets a human cytomegalovirus reactivation determinant": Supplmental Table 1

**SUPPORTING INFORMATION**

**SI Table 1. Candidate UL136 interactors from Y2H screen**

| **Name** | | **Known Functions** | **Accession Number** |
| --- | --- | --- | --- |
| **MYLIP** | Myosin regulatory light chain interacting protein | E3 Ubiquitin Ligase that mediates ubiquitination and degradation of myosin regulatory light chain (MRLC) | [XM_005249032.2](http://www.ncbi.nlm.nih.gov/nucleotide/767939837?report=genbank&log$=nucltop&blast_rank=2&RID=HXM24XAT013) |
| **PIK3R4** | Phosphoinositide-3-kinase, regulatory subunit 4 | Also called VPS15; serine-threonine protein kinase found at the cytoplasmic face of late Golgi and may regulate membrane trafficking in the late endocytic pathway | NM_014202.2 |
| **AP3B1** | Adapter-related protein complex 3, beta 1 subunit | Subunit of non-clathrin- and clathrin- associated adaptor protein complex 3 that functions in transmembrane protein sorting to lysosome or lysosomal-related organelles. | [NM_003664.4](http://www.ncbi.nlm.nih.gov/nucleotide/419636270?report=genbank&log$=nucltop&blast_rank=3&RID=TYYTT8R6014) |
| **CTSL2** | Cathespin L2 | Member of peptidase C1 family; lysosomal cysteine proteinase | [NM_001333.3](http://www.ncbi.nlm.nih.gov/nucleotide/320118899?report=genbank&log$=nucltop&blast_rank=1&RID=HXKG3K4Z013) |
| **TNFR-SF11B** | Tumor necrosis factor receptor superfamily, member 11B | TNFR superfamily member; may act as a decoy receptor for TRAIL thereby protecting against apoptosis | [NM_002546.3](http://www.ncbi.nlm.nih.gov/nucleotide/148743792?report=genbank&log$=nucltop&blast_rank=1&RID=HXKNSK2S013) |
| **CTNNAL1** | Catenin (cadherin-associated protein), alpha-like 1 | May modulate Rho pathway and act as anchor for PKA | [XM_011519161.1](http://www.ncbi.nlm.nih.gov/nucleotide/767958634?report=genbank&log$=nucltop&blast_rank=5&RID=HXKR6PMN016) |
| **MARK3** | MAP/microtubule affinity-regulating kinase 3 | Activated by phosphorylation, then phosphorylates MAP2, MAP4, and CDC25 | [XM_006720146.2](http://www.ncbi.nlm.nih.gov/nucleotide/767980561?report=genbank&log$=nucltop&blast_rank=7&RID=HXKV9Y92013) |
| **ZFP42** | Zinc finger protein 42 | Also called Rex1; marker of pluripotency, regulation is critical to maintaining pluripotent state in ES cells and may be involved in transcriptional regulation | [NM_001304358.1](http://www.ncbi.nlm.nih.gov/nucleotide/748983118?report=genbank&log$=nucltop&blast_rank=5&RID=HXKX1XKP016) |
| **MYLIP** | Myosin regulatory light chain interacting protein | E3 Ubiquitin Ligase that mediates ubiqutination and degradation of myosin regulatory light chain (MRLC) | [XM_005249032.2](http://www.ncbi.nlm.nih.gov/nucleotide/767939837?report=genbank&log$=nucltop&blast_rank=2&RID=HXM24XAT013) |
| **SCEL** | Sciellin | Localizes to periphery of cells; is a precursor of the cornified envelope of terminally differentiated keratinocytes | [XM_011535281.1](http://www.ncbi.nlm.nih.gov/nucleotide/767978191?report=genbank&log$=nucltop&blast_rank=11&RID=HXM399RG016) |
| **CHCHD4** | Coiled-coil-helix-coiled-coil-helix domain containing 4 | Mitochondrial chaperone to catalyze disulfide bonds. Required for the import and folding of small cysteine containing proteins into the mitochondrial intermembrane space | [NM_001098502.1](http://www.ncbi.nlm.nih.gov/nucleotide/148612858?report=genbank&log$=nucltop&blast_rank=2&RID=HXM4WX96016) |
| **ADSL** | Adenylosuccinate lyase | De novo synthesis of purines; catalyzes reactions in synthesis of AMP | [XM_011529976.1](http://www.ncbi.nlm.nih.gov/nucleotide/768023810?report=genbank&log$=nucltop&blast_rank=9&RID=HXMB68GG013) |
| **BTF3** | Basic transcription factor 3 | Forms stable complex with RNA polymerase II; required for transcription initiation | [NM_001037637.1](http://www.ncbi.nlm.nih.gov/nucleotide/83641884?report=genbank&log$=nucltop&blast_rank=1&RID=HXMC56XD016) |
| **DARS** | Aspartyl-tRNA synthetase | Part of multienzyme complex of aminoacyl-tRNA synthetases; charges its cognate tRNA with aspartate | [NM_001349.3](http://www.ncbi.nlm.nih.gov/nucleotide/648216370?report=genbank&log$=nucltop&blast_rank=2&RID=HXME7C2N016) |
| **TERF2IP** | Telomeric repeat binding factor 2, interacting protein | Part of a complex involved in telomere length regulation | [NM_018975.3](http://www.ncbi.nlm.nih.gov/nucleotide/294979143?report=genbank&log$=nucltop&blast_rank=2&RID=HXMF4226013) |
| **CCDC15** | Coiled-coil domain containing 15 | ? | [NM_025004.2](http://www.ncbi.nlm.nih.gov/nucleotide/144922719?report=genbank&log$=nucltop&blast_rank=1&RID=HXMGC6JZ016) |
| **CPORF43** | Chromosome 9, open reading frame 43 | ? | [NM_152786.2](http://www.ncbi.nlm.nih.gov/nucleotide/520262278?report=genbank&log$=nucltop&blast_rank=7&RID=HXMH8YZW013) |
